# Determining the neuronal ensembles underlying sex-specific social impairments following adolescent intermittent ethanol exposure

**DOI:** 10.1101/2023.03.21.533653

**Authors:** Trevor T. Towner, Matt A. Goyden, Harper J. Coleman, Mary K. Drumm, Isabella P. Ritchie, Kayla R. Lieb, Elena I. Varlinskaya, David F. Werner

## Abstract

Binge drinking during adolescence can have behavioral and neurobiological consequences. We have previously found that adolescent intermittent ethanol (AIE) exposure produces a sex-specific social impairment in rats. The prelimbic cortex (PrL) regulates social behavior, and alterations within the PrL resulting from AIE may contribute to social impairments. The current study sought to determine whether AIE-induced PrL dysfunction underlies social deficits in adulthood. We first examined social stimulus-induced neuronal activation of the PrL and several other regions of interest implicated in social behavior. Male and female cFos-LacZ rats were exposed to water (control) or ethanol (4 g/kg, 25% v/v) via intragastric gavage every other day between postnatal day (P) 25 and 45 (total 11 exposures). Since cFos-LacZ rats express β-galactosidase (β-gal) as a proxy for cFos, activated cells that express of β-gal can be inactivated by Daun02. β-gal expression in most ROIs was elevated in socially tested adult rats relative to home cage controls, regardless of sex. However, differences in social stimulus-induced β-gal expression between controls and AIE-exposed rats was evident only in the PrL of males. A separate cohort underwent PrL cannulation surgery in adulthood and were subjected to Daun02-induced inactivation. Inactivation of PrL ensembles previously activated by a social stimulus led to a reduction of social behavior in control males, with no changes evident in AIE-exposed males or females. These findings highlight the role of the PrL in male social behavior and suggest an AIE-associated dysfunction of the PrL may contribute to social deficits following adolescent ethanol exposure.

## 1. Introduction

Over 10% of the United States population 12 years of age and older are estimated to meet criteria for an alcohol use disorder (AUD) (Substance Abuse and Mental Health Services, SAMHSA, 2022). The early initiation of drinking, often reported during adolescence, is a strong predictor of developing AUD later in life (DeWit et al., 2000). Furthermore, early initiation of alcohol consumption promotes increased problematic/heavy alcohol use such as binge drinking (Aiken et al., 2018; Englund et al., 2008). Binge drinking is a characteristic drinking pattern of adolescents, with approximately 8% of those between the age of 12 and 20 years reporting at least one binge episode in the previous month (SAMHSA, 2022).

Adolescence is a sensitive developmental period (Larsen & Luna, 2018) during which the brain undergoes substantial remodeling. Alcohol consumption during this period can disrupt the natural maturation of the brain and potentially lead to behavioral/neurobiological deficits (Spear, 2018). The short-term consequences of adolescent binge drinking are numerous, with increases in risk taking (Marcotte et al., 2012; Stickley et al., 2013), decreases in attention (Squeglia et al., 2009), attenuated cognitive processing (Gil-Hernandez & Garcia-Moreno, 2016; Lees et al., 2020), and disrupted sleep (Ehlers et al., 2018) commonly reported. Along with these behavioral and cognitive deficits, some studies have found changes in brain structure and function such as altered grey matter volume (Perez-Garcia et al., 2022) as well as functional/structural connectivity (Correas et al., 2016; Morales et al., 2020; Sousa et al., 2019). However, the long-term consequences of binge drinking during adolescence have been difficult to measure given methodological limitations.

Using animal models, several long-term behavioral impairments resulting from adolescent intermittent ethanol (AIE) exposure have been identified (Crews et al., 2019). For example, AIE can lead to heightened anxiety-like behavior (Pandey et al., 2015; Sakharkar et al., 2019; Varlinskaya et al., 2020), increased risk-taking (Clark et al., 2012; Kruse et al., 2017), and impaired cognitive performance (Fernandez & Savage, 2017; Macht et al., 2020; Reitz et al., 2021) in adulthood. Several neurobiological changes have also been noted following AIE, including but not limited to increased neuroimmune signaling (Macht et al., 2020; Vetreno & Crews, 2018), increased blood brain barrier permeability (Vore et al., 2022), and altered glutamatergic functioning (Healey et al., 2021; Kasten et al., 2020). Behavioral deficits after AIE can be reversed through anti-inflammatory drugs and other pharmacological compounds acting on the neural systems affected be AIE (Macht et al., 2021; Swartzwelder et al., 2019; Vetreno et al., 2019), supporting a functional role of these findings.

Prior work from our lab has found that AIE consistently results in sex-specific social deficits evident in adulthood (Dannenhoffer et al., 2018; Towner et al., 2022a; Varlinskaya et al., 2020; Varlinskaya et al., 2017; Varlinskaya et al., 2014). Specifically, males exposed to ethanol during adolescence show decreased social preference and social investigation, with these findings absent in females. Furthermore, these male specific social deficits may increase the propensity for AIE-exposed males to consume ethanol when in a social environment (Towner et al., 2022b). However, our understanding of the neural mechanisms regulating AIE-induced social deficits are still limited.

Social behavior is complex and is characterized through multiple unique behaviors, all of which are likely regulated through specific neural mechanisms. The social brain network consists of several brain regions implicated in regulating various social behaviors (Kennedy & Adolphs, 2012; Newman, 1999). Although the critical role of the medial prefrontal cortex (mPFC) in regulation of social behavior is well established (Amodio & Frith, 2006; Kietzman & Gourley, 2023; Yizhar & Levy, 2021) multiple brain regions work jointly to control and modulate social behavior. Previous studies using circuit level approaches further support connectivity between frontal cortical regions and limbic structures (Huang et al., 2020; Iura et al., 2020; Li et al., 2021; Murugan et al., 2017) in regulating social behaviors.

We recently utilized cFos-LacZ transgenic rats that co-expressed beta-galactosidase (β-gal and cFos for assessment of neuronal activation of brain regions within the social brain network in animals previously exposed to AIE (Towner et al., 2022a). We found that water-exposed control males had positive relationships between social investigation and neuronal activation of the mPFC, consisting of the prelimbic (PrL) and infralimbic (IL) cortices. These relationships were absent in males exposed to AIE, findings that suggest that AIE-induced alterations of the mPFC may underly social impairments in these animals. Previously reported AIE-induced alterations within the mPFC include changes to gamma-aminobutyric acid (GABA) receptor mediated inhibitory signaling (Centanni et al., 2017; Rice et al., 2019; Trantham-Davidson et al., 2017), increased perineuronal net expression (Dannenhoffer et al., 2022), reduced astrocyte function (Healey et al., 2022), and dysregulated dopamine transmission (Trantham-Davidson et al., 2017). Therefore, it is possible that AIE-induced disruptions of mPFC maturation contributes to aberrant signaling in this region resulting in social deficits.

The current study was designed to determine whether AIE-associated changes in the PrL contribute to social impairment in AIE exposed males using cFos-LacZ rats. These rats express β-gal in cells recently activated by a certain stimulus, therefore allowing for pharmacogenetic inactivation of the selected neuronal ensembles with Daun02 to test whether these ensembles are directly involved in a behavioral response to this stimulus. In Experiment 1, we assessed the impact of AIE on neuronal activation within the mPFC and other brain regions induced by interaction with a same-sex social partner. In Experiment 2, we tested whether Daun02 inactivation of PrL neuronal ensembles previously activated by social interaction was sufficient to reverse AIE-induced social deficits in adult males.

## 2. General Method

### 2.1. Subjects

Male and female cFos-LacZ transgenic rats backcrossed onto Envigo-derived Sprague-Dawley background were bred and reared in our colony at Binghamton University for use in the current studies. cFos-LacZ breeders were obtained from the transgenic line originally developed by Dr. Curran while at Roche Institute of Molecular Biology. In these rats, the LacZ transgene that encodes for the enzyme β-gal is under the control of a cFos promoter, allowing for simultaneous expression of β-gal and cFos within the same cell (Fig. 1A). Male LacZ positive (LacZ+) rats were bred with wildtype females to produce litters with approximately 50% of offspring containing the LacZ gene. All litters were weaned on postnatal day (P) 21, at which offspring were group-housed with same-sex littermates (3-6 per cage). To confirm the presence of the LacZ gene, ear punches were taken between P21-24 and genotyped by TransnetYX (Cordova, TN). Once genotypes were determined and prior to P25, number of animals per cage was reduced to 3-4 subjects with variable genotype ratios present. Animals were maintained in a temperature (20-22° C), humidity, and light controlled (12 hours on/off, lights on at 0700h) vivarium. Water and food were provided *ad libitum*. Maintenance and experimentation were conducted in accordance with the animal use and care guidelines established by the National Institutes of Health and using protocols approved by Binghamton University Institutional Animal Care and Use Committee.

**Figure 1.**
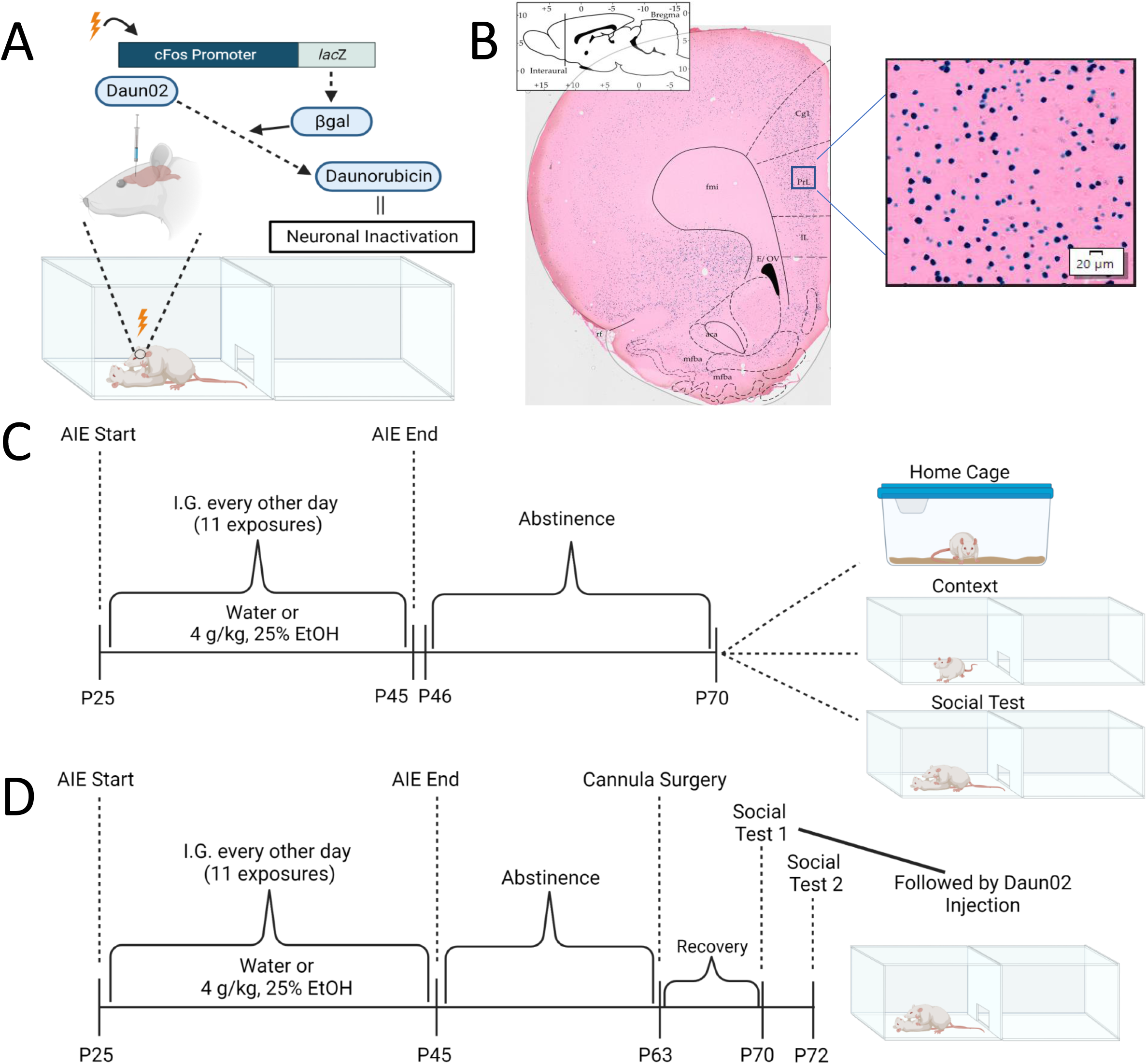
The cFos-LacZ transgenic rat model, adolescent intermittent ethanol (AIE) exposure, and social testing in adulthood. (A) cFos-LacZ rats have a LacZ gene on a cFos promoter and express β-galactosidase (β-gal) in recently activated neurons. The inert prodrug Daun02 is converted into daunorubicin by β-gal and leads to genetic interference in these cells ultimately inactivating them. (B) Representative image of β-gal labeling (indigo blue) in coronal slice. (C) Timeline of Experiment 1: AIE or water exposure and behavioral testing in adulthood. (D) Timeline of Experiment 2: AIE or water exposure followed by cannulation surgery and social testing in adulthood. Scale bar, 20µm.

### 2.2. Adolescent Intermittent Ethanol (AIE) Exposure

Rats were exposed to water (control) or ethanol (AIE) via intragastric gavage every other day from P25-45 for a total of 11 exposures (see Figure 1C-D for timeline). Ethanol was administered at 4 g/kg (25% v/v in tap water), and controls received an isovolumetric gavage with tap water. Previous studies from our lab have found this regimen to produce blood ethanol concentrations between 200 mg/dL at the start and 125 mg/dL at the end of the AIE exposure period (Kim et al., 2019). Exposures were conducted between 1200-1600h by a trained experimenter to minimize stress associated with the gavage procedure. All rats in a cage were given an identical exposure, regardless of genotype, however only LacZ+ rats were used for the current experiments. Following the final exposure (P45), rats were left undisturbed in their cages until adulthood (∼P70 Experiment 1, ∼P63 Experiment 2).

### 2.3. Modified Social Interaction Test

Social behavior was assessed using the modified social interaction test in accordance with our previous studies (Dannenhoffer et al., 2018; Towner et al., 2022a; Varlinskaya et al., 2020), Upon reaching adulthood (∼P70), LacZ+ rats were weighed, marked for identification, and placed into a testing apparatus (46.0 x 30.3 x 30.5 cm) that consisted of two equally sized chambers separated by an aperture, allowing for movement between compartments. Each apparatus contained pine shavings at the bottom and were cleaned with 6% hydrogen peroxide between sessions. Testing was conducted in the presence of white noise and in dim lighting (10-15 lux). Experimental rats were allowed to habituate to the testing environment for 30 minutes prior to the introduction of a novel, same-sex social partner weighing between 10-30 grams less. Social behavior was video recorded for 10 minutes, however the animals remained together for a total of 60 minutes.

Two primary social behaviors were assessed during the 10-minute recorded session: social investigation and social preference/avoidance coefficient. Social investigation was quantified as the frequency of sniffing any part of the social partner. The social preference/avoidance coefficient was calculated using the following formula: (crossovers to the partner – crossovers away from the partner) / (total # of crossovers to and away from the partner) x 100. Positive values are generally associated with social preference and negative values associated with social avoidance. All behaviors were hand-scored by a trained observer that was blind to experimental condition.

### 2.4. Tissue Collection and Sectioning

Following the 60-minute social interaction period, experimental animals were anaesthetized with an intraperitoneal injection of sodium pentobarbital and transcardially perfused with 0.01M phosphate buffered saline (PBS) followed by 4% paraformaldehyde in 0.1M phosphate buffer (PB). Brains were extracted, post fixed in 4% paraformaldehyde for 90 minutes, and then placed into cryoprotectant (30% sucrose in 0.01M PBS) prior to flash-freezing in methyl-butane. A cryostat (CM1860, Leica Biosystems, Wetzlar, Germany) was used to generate 30 μm coronal sections that were collected into an antifreeze solution (30% ethylene glycol, 30% glycerol, 0.05M PB). Sections were maintained at -20° C until processed.

### 2.5. Enzymatic Histology

As a proxy for cFos, β-gal labeling was used as an indicator of neuronal activation. Every 6^th^ section containing each region of interest underwent X-gal enzymatic histology using methods adapted from previous literature (Koya et al., 2016; Towner et al., 2022a). First, slices were fixed in 0.1M PB supplemented with 5mM EGTA for a total of 15 minutes. Slices were then washed (twice) for 5 minutes in 0.1M PB. Next, slices were incubated overnight at 37° C in staining buffer (0.1M PB with 2mM MGCl_2_, 5mM potassium ferrocyanide, and 5mM potassium ferricyanide) with added x-gal stock (5-bromo-4-chloro-3-indolyl-β-D-galactoside in dimethylformamide, 1:50). Finally, slices underwent two 10-minute washes in 0.1M PB and were mounted on charged microscopy slides (Histobond, VWR, PA, US). Mounted slides were then dehydrated, counterstained using eosin to aid in visualization of β-gal labeling, and coverslipped (Fig. 1B).

### 2.6. Image Acquisition and Analysis

Slides were imaged using a VS200 slide scanner (Olympus, Center Valley, PA) equipped with a 10x magnification objective. Images were then processed in Halo (Indica Labs, Albuquerque, NM) through application of annotation layers into each region of interest. Neuronal activation was automatically quantified based on β-gal expression within each annotation layer using a user designed algorithm built around features such as indigo blue staining, soma size, and nuclear roundness. We have validated this automatic detection to have over 95% reliability with observer identified cells. Results from Halo were consolidated and exported for processing in analysis programs.

### 2.7. Experiment 1: neuronal activation induced by social interaction

#### 2.7.1 Experimental Design

The goal of this experiment was to assess the neuronal activation state of our selected regions of interest (ROIs) in relation to a social stimulus. To do so we used a 2 adolescent exposure (water, AIE) X 3 testing condition (home cage, context, social) X 2 sex (male, female) design, with n =10-14 LacZ+ animals per group. No more than two animals per sex per litter per condition were used to minimize the influence of litter on the results. A total of 144 LacZ+ animals were used. Three testing conditions were used which allowed us to determine neuronal activation states associated with baseline AIE-induced changes (home cage), with exposure to the testing context (context), and with exposure to the social stimulus (social). Animals in all test conditions were exposed during adolescence to water or ethanol, as described above, and allowed to age into adulthood. On P70, rats in the home cage condition were taken from their cage and immediately euthanized using transcardial perfusion (see section 2.4.). Rats in the context condition were weighed and placed into the testing apparatus for a total of 90 minutes prior to being perfused. Finally, rats in the socially tested group were placed alone in the apparatus for a 30-minute habituation period, and then a social partner was placed in the apparatus for a 60-miute social interaction period (a total of 90 minutes in the context) after which experimental subjects were perfused.

All brains were extracted/processed in an identical fashion (see section 2.4.). Neuronal activation level was determined through quantification of β-gal labeling across several brain regions previously shown to be involved in social behavior (Kennedy & Adolphs, 2012; Ko, 2017). A total of 9 ROIs related to the social brain network were assessed: ventral orbitofrontal cortex (vOFC), lateral OFC, PrL, IL, Nucleus accumbens core (NAcC), Nucleus accumbens shell (NAcSh), central amygdala (CeA), basolateral amygdala (BLA), medial amygdala (MeA) (see Table 1 for coordinates). Within each ROI, a 400 x 400 μm box was placed, and the number of β-gal+ cells counted, with final values extrapolated to a number/mm^2^ (see Fig. 1B for example).

**Table 1.** Anatomic location of each region of interest.

| **Region of Interest** | **Location**<br>**(in reference to Bregma)** |
| --- | --- |
| *Lateral Orbitofrontal (lOFC)* | 4.68 – 2.76 mm |
| *Ventral Orbitofrontal (vOFC)* | 4.68 – 3.00 mm |
| *Prelimbic cortex (PrL)* | 4.40 – 2.52 mm |
| *Infralimbic cortex (IL)* | 3.72 – 2.52 mm |
| *Nucleus Accumbens Core (NAcC)* | 2.76 – 1.08 mm |
| *Nucleus Accumbens Shell (NAcSh)* | 2.76 – 0.96 mm |
| *Central Amygdala (CeA)* | -1.56 - -3.24 mm |
| *Basolateral Amygdala (BLA)* | -1.72 - -3.36 mm |
| *Medial Amygdala (MeA)* | -1.72 - -3.36 mm |

Given our prior findings of the PrL neuronal activation state positively correlating with social investigation in control males (Towner et al., 2022a), our primary hypothesis was that AIE-exposed males would have lower neuronal activation than water-exposed control males following exposure to a social stimulus. We expected sex-specific effects of AIE, with females displaying no difference in neuronal activation of the PrL between water- and AIE-exposed groups. In addition, we hypothesized that animals in the social testing condition would have the greatest number of β-gal+ cells in the ROIs given their known contribution to social behavior.

#### 2.7.2. Statistical Analysis

We have previously found AIE-induced social alterations only in males (Dannenhoffer et al., 2018; Towner et al., 2022a; Varlinskaya et al., 2020), therefore analyses were conducted within each sex separately. Social behaviors (investigation and coefficient) were analyzed with independent samples t-tests between exposure conditions (water, AIE). Two-way analyses of variance (ANOVAs) were used to assess the number of β-gal+ cells in each ROI across the three testing conditions (home cage, context, social) between water and AIE, independently for each sex. Given our previous findings of social investigation frequency correlating with PrL β-gal expression in water-exposed control males and our specific hypothesis, we used a priori comparisons to determine differences in PrL activation between water- and AIE-exposed animals in the socially tested condition. In other ROIs, interactions were assessed with follow up comparisons with Bonferroni corrections, focused on differences between exposure groups within each testing condition. Subjects with β-gal expression beyond two standard deviations from the mean were considered outliers and removed from analyses. Final group sizes ranged between 9-14 (individual values and sample size depicted in each figure).

### 2.8. Experiment 2: Daun02-induced inactivation of the prelimbic cortex

#### 2.8.1. Experimental Design

To determine whether neuronal activation differences observed in Experiment 1 were functionally relevant to social deficits following AIE, Experiment 2 used Daun02 to silence neurons in the PrL (Fig. 1A). A 2 adolescent exposure (water, AIE) X 2 drug (vehicle, Daun02) X 2 sex (male, female) design was used, with n =13-14 LacZ+ animals per group (total N = 110). Subjects were exposed to water or AIE and allowed to age into adulthood. On P63, cannulae were bilaterally implanted in the PrL (see section 2.8.2.) and allowed one week to recover from surgery before social testing. Following surgery, animals were isolate housed for the remainder of the study. To induce neuronal activation associated with a social stimulus, rats were exposed to social partners on P70 for a total of 60 minutes. Procedures were identical to that of Experiment 1. Immediately following the 60-minute social interaction, Daun02 was microinjected (see section 2.8.3.) into the PrL cortex to inactivate the cells expressing β-gal. Rats were socially tested a second time on P72 to determine if inactivation of PrL neuronal ensembles by Daun02 altered subsequent social behavior. Again, rats were placed alone in the chamber for a 30-minute habituation period followed by a 60-minute social interaction. Subjects were perfused, brains extracted and post-fixed, as previously described. See figure 1D for a timeline of Experiment 2.

Brains were processed for β-gal expression as described above. Our primary focus was on the number of social stimulus associated β-gal+ cells in the PrL following administration of vehicle (5% Dimethyl Sulfoxide (DMSO), 5% Tween 80, 90% artificial cerebrospinal fluid (ACSF)) or Daun02. In addition, the expression of β-gal in the ROIs described in Experiment 1 was also quantified to determine whether altering neuronal activation of the PrL could affect neuronal activation of other ROIs implicated in social behavior.

Experiment 2 was guided by three hypotheses. First, we hypothesized that Daun02-injected animals would have lower β-gal expression in the PrL than vehicle-injected controls, regardless of sex. Second, it was expected that AIE-induced social deficits would be reversed through inactivation of PrL cells, and therefore no differences in social behavior would be evident between male water- and AIE-exposed groups. No effects of AIE on female social behavior were expected. Lastly, we hypothesized that inactivation of the PrL would increase neuronal signaling in the extended amygdala due to PrL top-down control.

#### 2.8.2. Cannulation Surgery

Following adolescent exposure, rats were left undisturbed in their cages until ∼P62. On P63, rats were deeply anesthetized with 3% isoflurane and placed into a stereotaxic frame. Once head-fixed, animals were maintained on 1-3% isoflurane for the remainder of the surgery. Bilateral holes were drilled into the skull using coordinates for the PrL (AP: +3.2mm, ML: ±0.75mm, and DV: -2.5mm). Bilateral, 26-gauge, steel-guide cannulae (PlasticsOne, Roanoke, VA) were slowly lowered until the pedestal base reached the skull. Additional holes were drilled for implanting screws. Dental cement was used to affix the cannulae in place. Dummy cannulae were inserted into the guide cannulae until time of microinjection. Rats regained consciousness in warmed cages. To minimize pain, intraperitoneal (IP) injections of buprenorphine HCl (0.03mg/kg) were administered immediately prior to surgery and every 12 hours for 48 hours. Rats were isolate housed for the remainder of the study and allowed a minimum of one week to recover prior to behavioral testing. During recovery, rats were handled, and dummy cannulae removed/reinserted to acclimate animals to manipulation of the headcap.

#### 2.8.3. Microinjection Procedure

Sixty minutes following introduction of the social partner on the first test day, experimental animals were taken to a separate room for a microinjection of either Daun02 (2μg/0.5 μL/side) or vehicle. Cannula caps and dummy cannulae were removed, and injectors extending 1mm past each guide cannula were lowered into the PrL. Each animal’s respective drug was microinjected over two minutes and injectors were left in place for an additional two minutes to prevent backflow during injector removal. Dummy cannulae were reinserted and animals were returned to the colony. The second behavioral test took place two days later after allowing for Daun02 to produce sufficient inactivation (de Guglielmo et al., 2016).

#### 2.8.4. Cannula Placement Verification

Accurate cannula placements were verified through enzymatic labeling of β-gal of every 6^th^ slice containing the PrL. Cannula tracks were identified in individual slices and ascribed approximate coordinates by an experimenter. Animals with cannula tracks outside of the PrL were excluded from further analysis (1 water-exposed male injected with vehicle, 1 water-exposed male injected withDaun02, and 1 water-exposed female injected with Daun02).

#### 2.8.5. Statistical Analysis

As in Experiment 1, data were separated by sex prior to analyses given our prior findings of AIE-induced sex-specific social deficits (Dannenhoffer et al., 2018; Towner et al., 2022a; Varlinskaya et al., 2014). Social behavior was assessed using a 2 adolescent exposure (water, AIE) X 2 drug (vehicle, Daun02) ANOVA separately for each sex. Similar two-way ANOVAs were conducted for analyzing β-gal expression in the PrL and other ROIs. Follow up comparisons with Bonferroni tests were conducted to further analyze significant interactions. Outliers were removed if outside of two standard deviations of the mean. Due to surgical complications, incorrect cannula placements, and outliers, final N per group was between 11 – 13.

## 3. Results

### 3.1. Experiment 1: neuronal activation induced by social interaction

#### 3.1.1. Social Behavior

An independent samples t-test of social investigation revealed a significant difference between water- and AIE-exposed males, *t* (23) = 3.686, *p* = 0.001, with AIE males demonstrating lower social investigation than water-exposed controls (Fig. 2A). However, no difference was found between water- and AIE-exposed males when assessing social preference/avoidance coefficient (Fig. 2B). Similarly, analyses of social investigation and social preference/avoidance coefficient in females revealed no significant differences between water- and AIE-exposed subjects (Fig. 2D-E).

**Figure 2.**
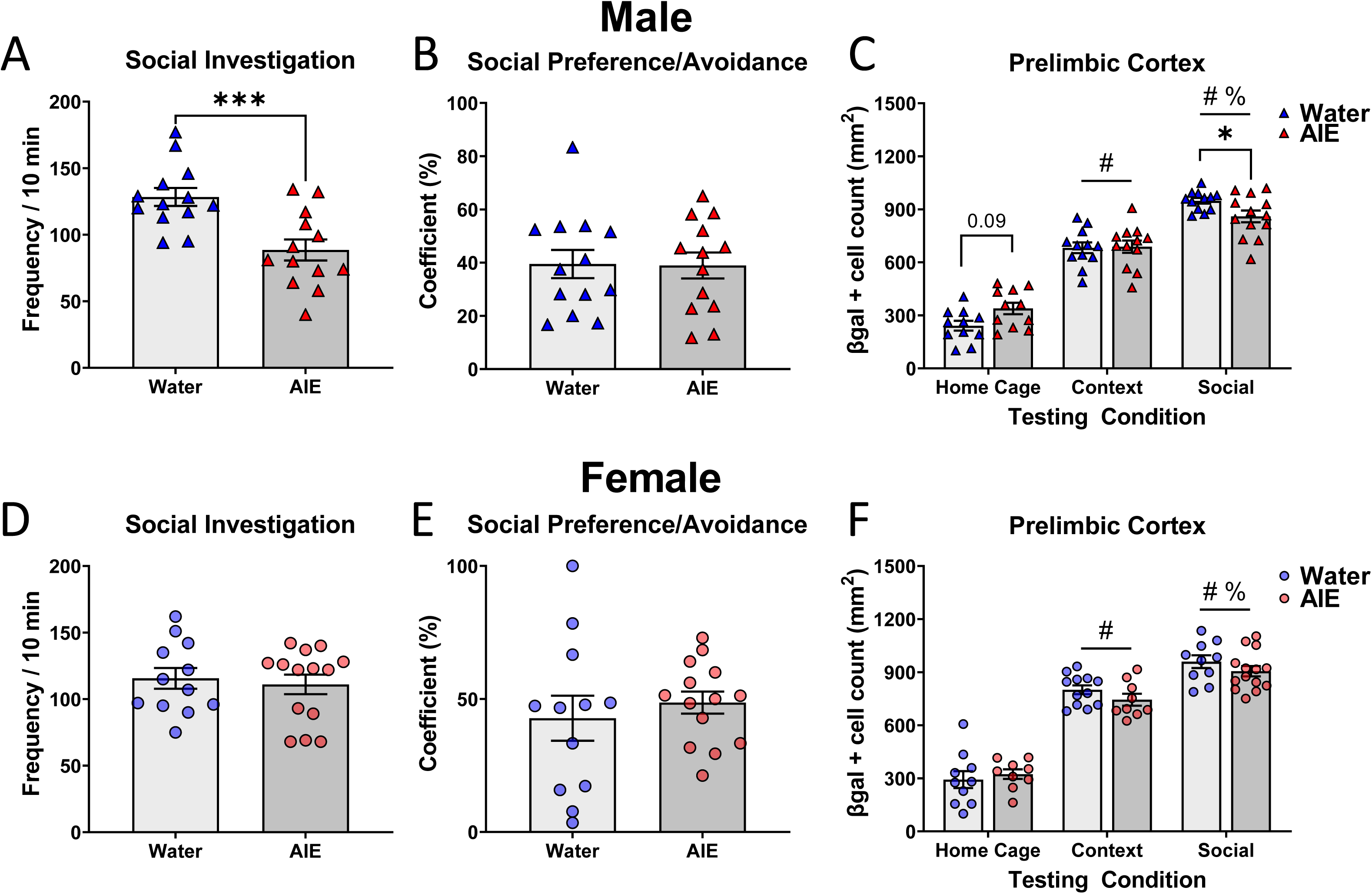
AIE-induced changes of social responding and neuronal activation in the PrL. Social investigation, social preference, and neuronal activation of the PrL indexed via β-gal+ cell counts in water- and AIE-exposed male (A, B, C, respectively) and female (D, E, F, respectively) rats taken from home cage, placed into the context, or exposed to a social partner. Significant differences between water and AIE exposure conditions are marked with asterisks (* *p* < 0.05 and *** *p* < 0.001); significant changes relative to the home cage condition are marked with number signs (# *p* < 0.001), significant changes relative to the context condition are marked with percent signs (% *p* < 0.001).

#### 3.1.2. Neuronal Activation (**β**-gal)

To determine the effects of AIE and testing condition on the neuronal activation across the ROIs (see Table 1), we conducted two-way (2 adolescent exposure x 3 testing condition) ANOVAs within each sex separately. In males, ANOVAs of β-gal expression revealed main effects of testing condition across all ROIs (for all F and p values see Table 2). Home cage animals had the lowest number of β-gal+ cells, with significant increases evident following placement in the testing context, and further significant increases in the number of β-gal+ cells seen in animals exposed to a social stimulus (Fig. 3A-B and Supplemental Fig. 1). However, in the CeA of males, significantly more β-gal+ cells were evident in the context and social test conditions relative to home cage β-gal expression (*p* < 0.05 and *p* < 0.001, respectively), with no difference seen between context and socially tested groups (*p* = 0.519, see Fig. 3C). In addition, the two-way (2 adolescent exposure x 3 testing condition) ANOVA of β-gal expression in the CeA revealed a main effect of adolescent exposure was identified, *F* (1, 65) = 4.102, *p* < 0.05. As seen in Figure 3C, males exposed to AIE had lower β-gal expression in the CeA than water-exposed counterparts, regardless of testing condition.

**Figure 3.**
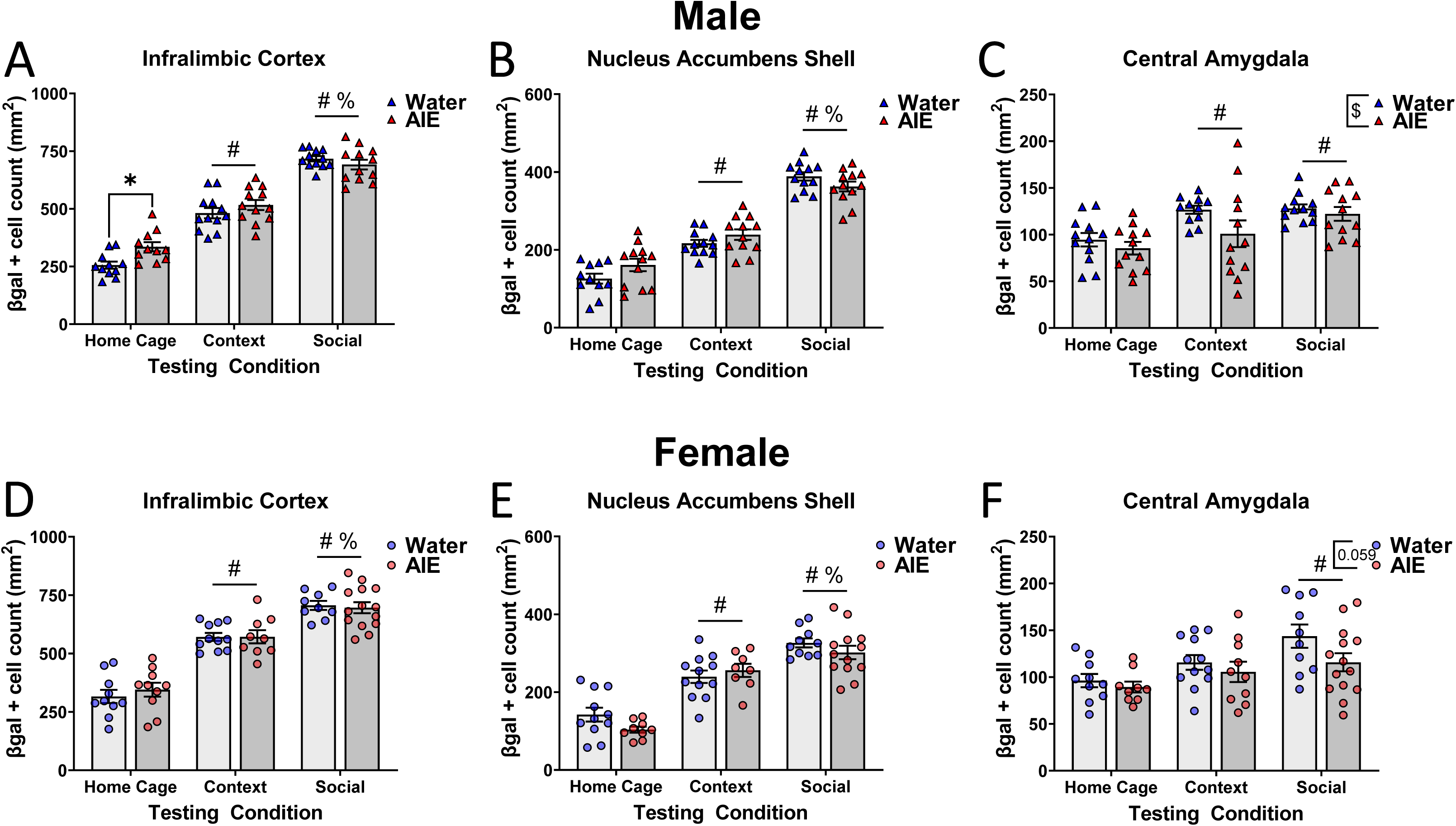
AIE-induced changes in neuronal activation of IL, NAcSh, and CeA. Neuronal activation of the IL, NAcSh, and CeA indexed via β-gal+ cell counts in water- and AIE-exposed male (A, B, C, respectively) and female (D, E, F, respectively) subjects taken from home, placed into the context, or exposed to a social partner. Significant differences between water and AIE exposure conditions are marked with asterisks (* *p* < 0.05); significant changes relative to the home cage condition are marked with number signs (# *p* < 0.001), significant changes relative to the context condition are marked with percent signs (% *p* < 0.001).

**Table 2.** Results of two-way ANOVAs of β-gal expression in each region of interest.

| Region of Interest | Results |  |  |
| --- | --- | --- | --- |
|  | Factor | Males | Females |
| *Lateral Orbitofrontal (lOFC)* | Adolescent Exposure | $F(1, 65) = 1.07, p = 0.31$ | $F(1, 58) = 0.25, p = 0.62$ |
|  | Testing Condition | **$F(2, 65) = 138.5, p < 0.0001$** | **$F(2, 58) = 111.7, p < 0.0001$** |
| | Exposure X Testing | $F(2, 65) = 1.99, p = 0.15$ | $F(2, 58) = 0.36, p = 0.70$ |
| *Ventral Orbitofrontal (vOFC)* | Adolescent Exposure | $F(1, 66) = 2.06, p = 0.16$ | $F(1, 57) = 0.76, p = 0.39$ |
|  | Testing Condition | **$F(2, 66) = 90.18, p < 0.0001$** | **$F(2, 57) = 104.7, p < 0.0001$** |
| | Exposure X Testing | $F(2, 66) = 2.85, p = 0.06$ | $F(2, 57) = 0.03, p = 0.97$ |
| *Prelimbic cortex (PrL)* | Adolescent Exposure | $F(1, 65) = 0.05, p = 0.83$ | $F(1, 58) = 0.89, p = 0.35$ |
|  | Testing Condition | **$F(2, 65) = 211.5, p < 0.0001$** | **$F(2, 58) = 179.5, p < 0.0001$** |
| | Exposure X Testing | **$F(2, 65) = 4.80, p = 0.01$** | $F(2, 58) = 1.02, p = 0.37$ |
| *Infralimbic cortex (IL)* | Adolescent Exposure | $F(1, 64) = 3.62, p = 0.06$ | $F(1, 57) = 0.10, p = 0.75$ |
|  | Testing Condition | **$F(2, 64) = 221.3, p < 0.0001$** | **$F(2, 57) = 113.4, p < 0.0001$** |
| | Exposure X Testing | **$F(2, 64) = 3.69, p = 0.03$** | $F(2, 57) = 0.32, p = 0.72$ |
| *Nucleus Accumbens Core (NAcC)* | Adolescent Exposure | $F(1, 67) = 0.73, p = 0.40$ | $F(1, 57) = 0.16, p = 0.69$ |
|  | Testing Condition | **$F(2, 67) = 46.93, p < 0.0001$** | **$F(2, 57) = 34.71, p < 0.0001$** |
| | Exposure X Testing | $F(2, 67) = 1.42, p = 0.25$ | $F(2, 57) = 0.69, p = 0.50$ |
| *Nucleus Accumbens Shell (NAcSh)* | Adolescent Exposure | $F(1, 65) = 1.06, p = 0.31$ | $F(1, 57) = 1.44, p = 0.23$ |
|  | Testing Condition | **$F(2, 65) = 174.7, p < 0.0001$** | **$F(2, 57) = 76.03, p < 0.0001$** |
| | Exposure X Testing | **$F(2, 65) = 3.30, p = 0.04$** | $F(2, 57) = 1.55, p = 0.22$ |
| *Central Amygdala (CeA)* | Adolescent Exposure | **$F(1, 65) = 4.10, p = 0.05$** | $F(1, 59) = 3.71, p = 0.06$ |
|  | Testing Condition | **$F(2, 65) = 9.686, p = 0.0002$** | **$F(2, 59) = 7.51, p = 0.001$** |
| | Exposure X Testing | $F(2, 65) = 0.84, p = 0.44$ | $F(2, 59) = 0.74, p = 0.48$ |
| *Basolateral Amygdala (BLA)* | Adolescent Exposure | $F(1, 65) = 1.16, p = 0.29$ | $F(1, 57) = 0.04, p = 0.85$ |
|  | Testing Condition | **$F(2, 65) = 88.23, p < 0.0001$** | **$F(2, 57) = 112.4, p < 0.0001$** |
| | Exposure X Testing | $F(2, 65) = 0.09, p = 0.91$ | $F(2, 57) = 0.39, p = 0.68$ |
| *Medial Amygdala (MeA)* | Adolescent Exposure | $F(1, 64) = 1.35, p = 0.25$ | $F(1, 57) = 0.41, p = 0.53$ |
|  | Testing Condition | **$F(2, 64) = 130.8, p < 0.0001$** | **$F(2, 57) = 73.05, p < 0.0001$** |
| | Exposure X Testing | $F(2, 64) = 2.97, p = 0.06$ | $F(2, 57) = 0.06, p = 0.94$ |

Bolded text indicates significant effect/interaction.

In the PrL of males, a two-way ANOVA of β-gal expression revealed an interaction of adolescent exposure and testing condition, *F* (2, 65) = 4.789, *p* = 0.01. Using a Bonferroni test, we found a trend toward a difference between water and AIE groups in the home cage condition (*p* = 0.089), however significance was not met. As seen in Figure 2C, in the home cage condition, AIE-exposed males tended to have more β-gal+ cells than controls, however significance was not met. Comparison of water- and AIE-exposed socially tested males revealed a significant difference (*p* < 0.05) with fewer β-gal+ cells activated by a social stimulus in males exposed to AIE than water-exposed counterparts. A significant interaction was also revealed for β-gal expression in the IL, *F* (2, 64) = 3.691, *p* < 0.05. Post hoc comparisons revealed a significant difference between adolescent exposures in home cage subjects (*p* < 0.05), with AIE-exposed males having greater β-gal expression than water-exposed males (Fig. 3A). In addition, a two-way ANOVA of β-gal expression in the NAcSh revealed an adolescent exposure by testing condition interaction, *F* (2, 65) = 3.298, *p* < 0.05. However, post hoc comparisons revealed no significant differences between adolescent exposure groups within each testing condition (Fig. 3B).

In females, two-way (2 adolescent exposure x 3 testing condition) ANOVAs revealed significant differences between testing conditions for all ROIs (Table 2). Generally, home cage females had low β-gal expression, with significantly more β-gal+ cells in the context females, and further increases in β-gal expression in socially tested females (Fig. 3D-E and Supplemental Fig. 2). However, in the CeA of females, post hoc analyses of this main effect identified only a significant increase in β-gal expression associated with social testing relative to the home cage condition (*p* < 0.001, Fig. 3F). The analysis of β-gal expression in the CeA also revealed a trend for a main effect of adolescent exposure was found, *F* (1, 59) = 3.710, *p* = 0.0589. Like males, AIE-exposed females tended to have a lower number of β-gal+ cells in the CeA than water-exposed controls, although this effect of AIE did not reach statistical significance (Fig. 3F). Two-way ANOVAs of other ROIs failed to reveal main effects of adolescent exposure or an interaction of adolescent exposure and testing condition (Table 2).

### 3.2. Experiment 2: Daun02-induced inactivation of the prelimbic cortex

Given that the PrL was the only region where we identified differences in neuronal activation following an interaction with a social stimulus between male adolescent exposure conditions, we targeted this region for neuronal inactivation using Daun02.

#### 3.2.1. Social Behavior

Two-way (2 adolescent exposure x 2 drug) ANOVAs were performed to determine the effect of Daun02-induced inactivation of PrL neuronal ensembles on social behavior for each sex independently. In males, analysis of social investigation revealed a significant adolescent exposure x drug interaction, *F* (1, 43) = 7.486, *p* < 0.01. As seen in Figure 4a, AIE-exposed males injected with vehicle had significantly lower social investigation than water-exposed counterparts (*p* = 0.01). In water-exposed males, inactivation of the PrL by Daun02 significantly reduced social investigation relative to vehicle-injected rats (*p* < 0.05), with no effect of Daun02 evident in AIE-exposed males (Fig. 4A). In contrast to social investigation, neither adolescent exposure nor Daun02 administration influenced social preference of male rats (Fig. 4B).

**Figure 4.**
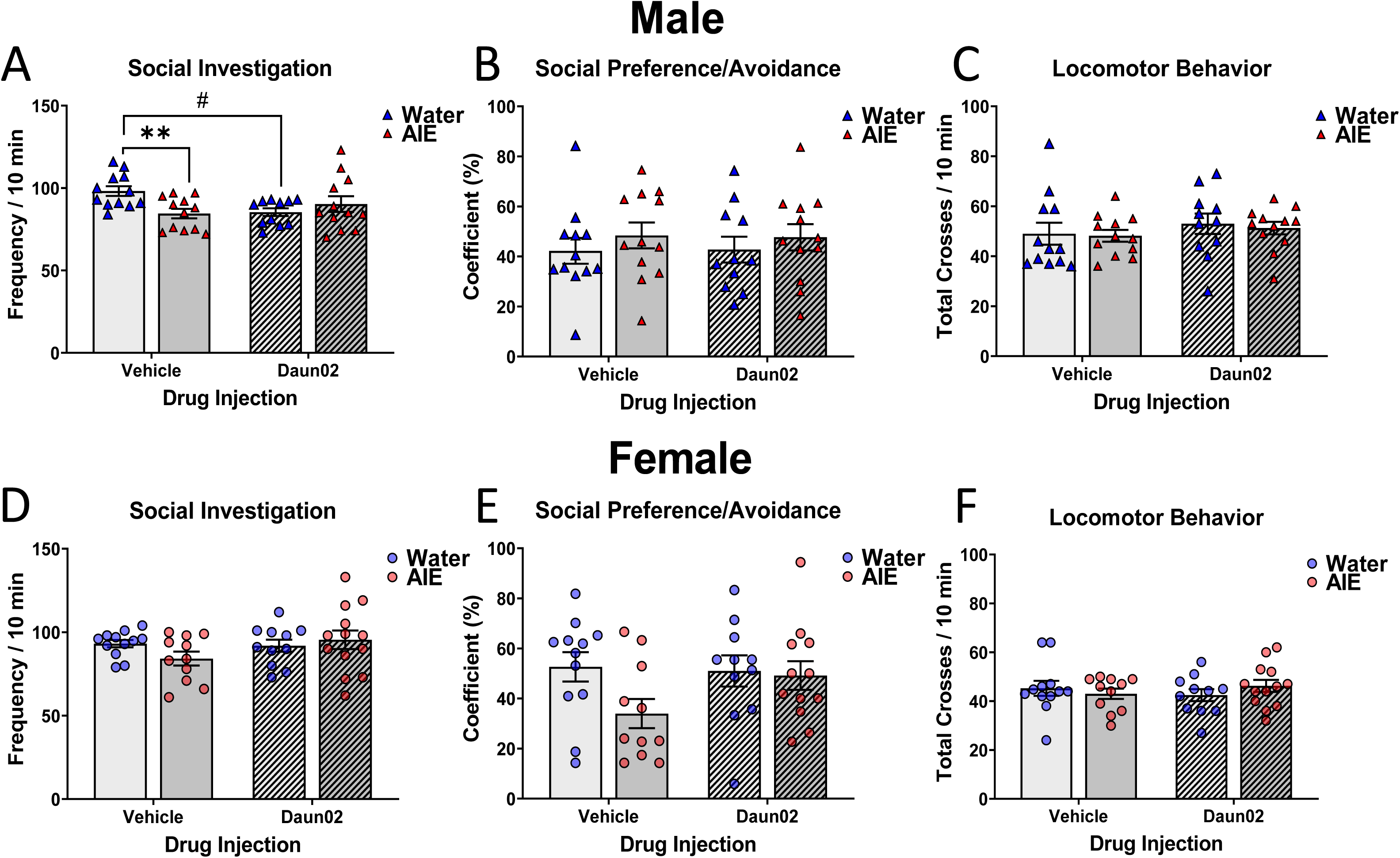
Daun02 effects on social behavior and locomotor activity. Social investigation, social preference/avoidance, and locomotor activity of water- and AIE-exposed male (A, B, C, respectively) and female (D, E, F, respectively) rats. Significant differences between water and AIE exposure conditions are marked with asterisks (** *p* < 0.01) AIE vs water-exposed controls; significant differences between vehicle and Daun02 are marked with number signs (# *p* < 0.05).

In females, two-way (2 adolescent exposure x 2 drug) ANOVAs of social investigation and social preference did not find significant main effects or interactions (Fig. 4D-E).

To control for potential locomotor differences leading to the observed Daun02 effect in water-exposed males, we conducted two-way ANOVAs on the total number of crossovers (movements between compartments) made during the test session. Analyses revealed no differences in locomotor activity (total crossovers made during the social interaction) were for either males or females (Fig. 4C & F).

#### 3.2.2. Neuronal Activation

Prior to analysis, we evaluated the placement of cannula in the PrL through visualization of cannula tracks within the region (Fig. 5A). Due to a lack of correct placement, three animals were removed from all analyses. Representative images of β-gal expression following either injection with vehicle or Daun02 can be seen in Figure 5B-C.

**Figure 5.**
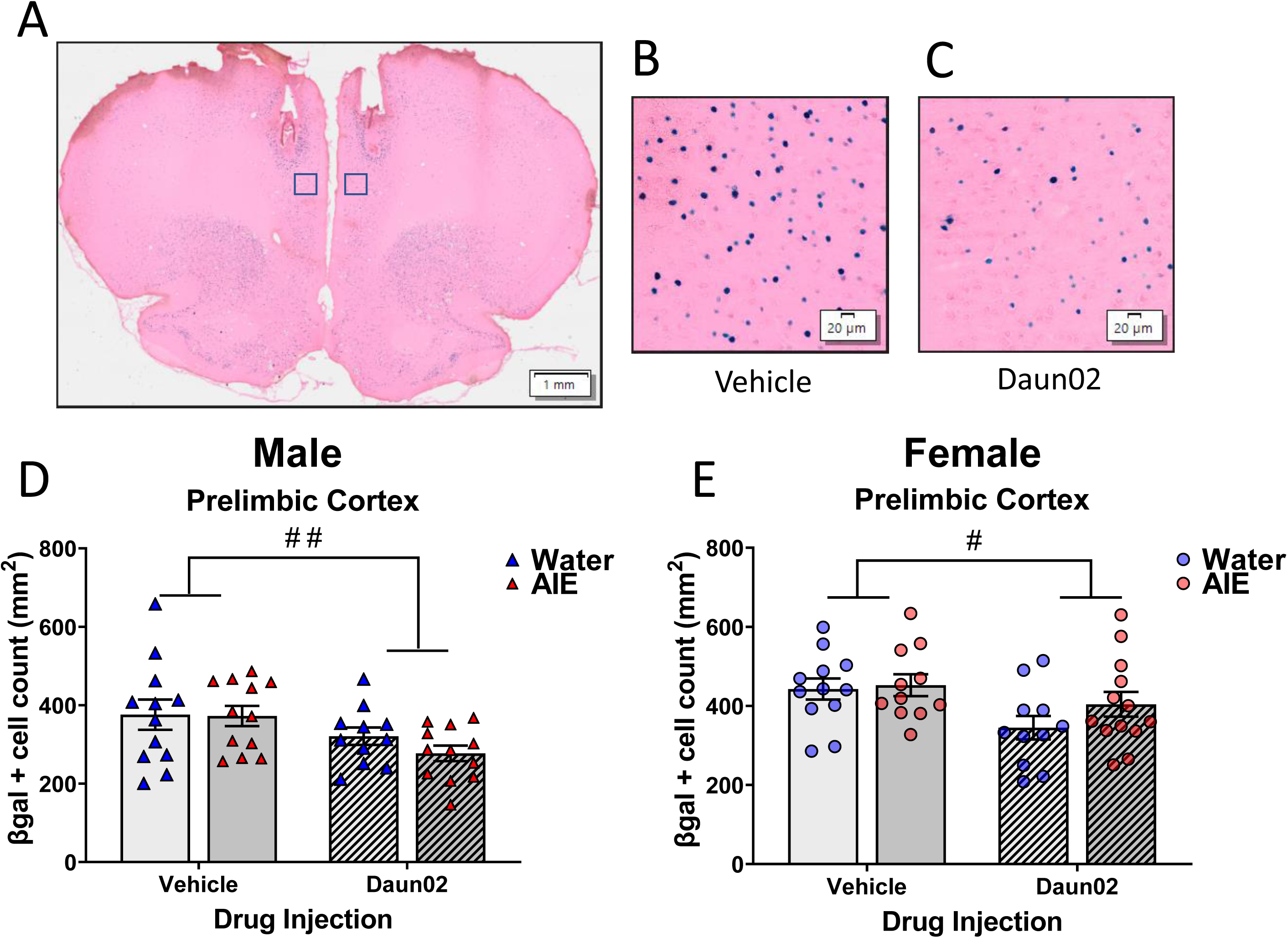
Daun02 effects on neuronal activation of the PrL indexed via β-gal expression. Representative image of PrL cannula placement (A) scale bar 1mm. Example images from the PrL of vehicle- (B) and Daun02-injected (C) subjects, scale bar 20µm. β-gal+ cell counts in the PrL of males (D) and females (E). Significant differences between vehicle and Daun02 are marked with number signs (# *p* < 0.05, ## *p* < 0.01).

The effectiveness of Daun02 to inactivate neuronal ensembles within the PrL was assessed using a two-way ANOVA of PrL β-gal+ cells elicited during the second social interaction. In males, a main effect of drug (vehicle; Daun02) was revealed, *F* (1, 43) = 7.282, *p* < 0.01, with Daun02 injected rats having fewer β-gal+ cells in the PrL (Fig. 5B-D). Similarly, a two-way ANOVA of β-gal+ cells in female PrL revealed a main effect of drug, *F* (1, 43) = 6.292, *p* < 0.05. As seen in Figure 5E, females injected with Daun02 had a lower number of cells expressing β-gal than vehicle injected counterparts.

To understand how PrL inactivation influenced downstream circuitry, we examined β-gal expression in other ROIs that are known to be directly/indirectly regulated by PrL signaling. In males, two-way ANOVAs revealed main effects of drug on β-gal expression in the NAcC, *F* (1, 43) = 5.689, *p* < 0.05, NAcSh, *F* (1, 43) = 7.816, *p* < 0.01, and CeA, *F* (1, 43) = 11.15, *p* < 0.01. All three regions displayed significantly lower number of β-gal+ cells in males that received Daun02 injections compared to vehicle injected controls (Fig. 6A-C). Two-way ANOVAs of β-gal expression in the same regions of females identified main effects of drug in the NAcC, *F* (1, 43) = 5.516, *p* < 0.024, and CeA, *F* (1, 43) = 5.30, *p* < 0.05. However, in the NAcSh, only a trend toward a main effect of drug was found, *F* (1, 43) = 3.612, *p* = 0.064. As was seen in males, females injected with Daun02 into the PrL tended to have a lower number of β-gal+ cells in the NAcC, NAcSh, and CeA (Fig. 6D-F).

**Figure 6.**
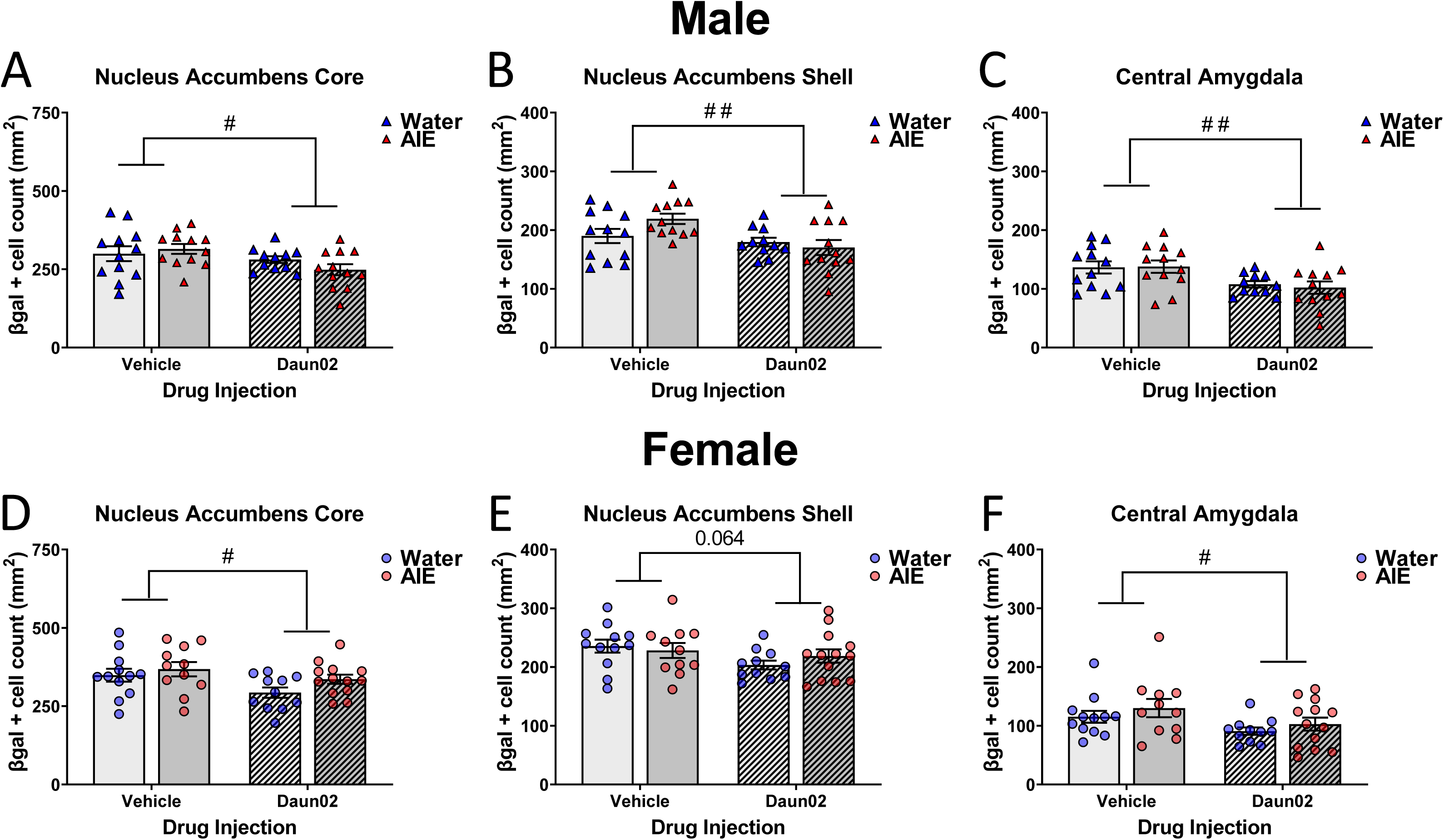
Neuronal activation of subcortical brain regions following Daun02 injection in the PrL. Neuronal activation indexed via β-gal expression of NAcC, NAcSh, and CeA in water- and AIE-exposed males (A, B, C, respectively) and females (D, E, F, respectively). Significant differences between vehicle and Daun02 are marked with number signs (# *p* < 0.05, ## *p* < 0.01).

## 4. Discussion

The current study was designed to identify neuronal ensembles that contribute to AIE-induced sex-specific social impairments. We previously found that males exposed to ethanol during adolescence display social deficits in adulthood, with these deficits possibly associated with a disruption of PrL neuronal activity (Towner et al., 2022a). However, due to the limited scope of our previous work, we were unable to functionally connect PrL neuronal ensembles activated during social interaction with AIE-induced social impairments. To further our understanding of PrL involvement in social deficits following AIE, we assessed β-gal expression, as a proxy of cFos, across three testing conditions, including home cage condition that allowed assessment of baseline differences between animals previously exposed to water or AIE. At baseline (home cage condition), AIE-exposed males had a greater number of β-gal+ neurons in the PrL, further supporting a potential dysregulation of localized excitatory/inhibitory signaling balance in the PrL identified by others (Centanni et al., 2017; Galaj et al., 2020; Salling et al., 2018; Trantham-Davidson et al., 2017).

The mPFC, including both the PrL and IL, undergoes substantial maturation and refinement during adolescence (Larsen & Luna, 2018). Thus, it is not surprising that the mPFC of adolescents is uniquely vulnerable to ethanol. Previous research has identified several AIE-associated changes in excitatory (Galaj et al., 2020; Salling et al., 2018), inhibitory (Centanni et al., 2017), cholinergic (Fernandez & Savage, 2017), and dopaminergic (Obray et al., 2022; Shnitko et al., 2016; Trantham-Davidson et al., 2017) signaling in the mPFC. Increased β-gal expression in the PrL and IL of AIE-exposed males in the home cage condition may be a result of a blunted excitatory/inhibitory remodeling that typically occurs during adolescence (Caballero et al., 2021). During adolescence, tonic GABA_A_-mediated current in PrL pyramidal neurons increases, and AIE reduces this developmental rise of inhibitory signaling through alterations of δ containing GABA_A_ receptor efficacy (Centanni et al., 2017). In addition, PrL fast-spiking interneurons display reduced intrinsic excitability as well as lower glutamate receptor dependent currents following AIE (Trantham-Davidson et al., 2017). The reduced inhibitory signaling in the PrL following AIE may lead to heightened basal activation as evident in current study, with this dysregulation in activity potentially making animals more prone to behavioral impairments.

The role of the mPFC in regulating social behavior has been extensively studied (Amodio & Frith, 2006; Yizhar & Levy, 2021). For example, in humans with autism spectrum disorder, a psychiatric disorder often characterized by impaired social behaviors, the mPFC is found to have abnormal development of cortical layers and specific expression of neuronal phenotypes (Stoner et al., 2014; Varghese et al., 2017) as well as reduced regional connectivity (Kana et al., 2009; Wass, 2011). Disruption of the excitatory/inhibitory balance in the mPFC is implicated in autism as well (Bicks et al., 2015; Canitano & Palumbi, 2021; Ferguson & Gao, 2018a; Nelson & Valakh, 2015), with supporting evidence from rodent studies (Ferguson & Gao, 2018b; Selimbeyoglu et al., 2017; Yizhar et al., 2011). It is posited that a shift toward hyperexcitability in the mPFC promotes social deficits in animal models of autism (Bicks et al., 2015). The basal increase in mPFC β-gal expression observed in AIE-exposed males in the current study may represent this hyperexcitable state.

Interestingly, the number of β-gal+ cells identified in socially tested subjects was significantly lower in AIE-exposed males, suggesting reduced neuronal activation of the PrL in animals that display social deficits. This finding is in opposition to that of home cage subjects, and may be a failure of AIE-exposed males to properly recruit neuronal ensembles necessary to display levels of social investigation similar to water-exposed controls. However, the finding of lower β-gal expression and the interpretation of lower activation in the PrL being associated with social deficits contradicts others who found elevated PrL excitation to result in social impairments (Ferguson & Gao, 2018b; Kim et al., 2020; Selimbeyoglu et al., 2017; Yizhar et al., 2011). For example, optogenetic stimulation of GABAergic signaling or inhibition of pyramidal neurons rescues social deficits in Contactin associated protein like 2 knockout mice (Selimbeyoglu et al., 2017), although knockout of this gene product results in hyperactivity and epileptic seizures (Penagarikano et al., 2011) that are not present following AIE. Alternatively, several studies have identified social impairments being a result of hypoexcitability of the mPFC, often associated with lower pyramidal neuron firing (Asgarihafshejani et al., 2019; Brumback et al., 2018; Dani et al., 2005; Howell et al., 2017; Sacai et al., 2020; Sceniak et al., 2016).

Furthermore, increasing inhibitory signaling using chemogenetics, thus similarly reducing overall activity in this region, reduces social preference (Ferguson & Gao, 2018b). Taken together with these findings, our results suggest that attenuation of PrL responding to a social stimulus, indexed via reduced neuronal activation, may be associated with the social deficits evident in AIE-exposed males.

In contrast to the PrL, IL β-gal expression elicited by social interaction was not affected by AIE. Others have identified opposing roles of the PrL and IL in fear expression and extinction (Sierra-Mercado et al., 2011), therefore it is not surprising that these regions may uniquely contribute to social behavior. In fact, evidence supports a differential role of these regions in regulating social behavior (Abe et al., 2019; Huang et al., 2020; Kim et al., 2015), suggesting that these regions may not contribute equally to AIE-induced reductions of social investigation.

In addition to AIE-induced changes of mPFC activation state, the CeA of both sexes was found to have reduced neuronal activation (β-gal) after AIE exposure, regardless of testing condition. It has previously been shown that AIE exposure reduces activity-regulated cytoskeletal (Arc) gene expression (Kyzar 2019; Pandey 2015), decreases brain derived neurotrophic factor (BDNF) (Pandey 2015), and lowers neuropeptide Y protein (Kokare 2017) in the CeA, with these alterations associated with various behavioral impairments such as anxiety-like behavior (Kokare et al., 2017; Kyzar et al., 2019; Kyzar et al., 2017; Pandey et al., 2015). In our previous study, reduced neuronal activation of the CeA was evident in AIE-exposed males, whereas females displayed a moderate increase in CeA neuronal activation (Towner et al., 2022a). These contradicting findings clearly show that more work is needed to understand the long-term consequences of AIE on CeA neuronal activation between sexes. Our finding of altered CeA β-gal expression in all testing conditions suggests that AIE-induced changes in CeA neuronal activation do not contribute to social behavior. To date, limited evidence of CeA involvement in social investigation has been identified in laboratory rodents although it may have a role in other social behaviors such as aggression (Veenema & Neumann, 2008) and social fear (Andraka et al., 2021). However, given the CeA’s prominent role in anxiety (Gilpin et al., 2015), it is likely that the observed disruptions in CeA neuronal activation contribute to AIE-induced non-social anxiety-like behavioral alterations as we (Varlinskaya et al., 2020) and others have previously observed (Crews et al., 2019).

For further understanding the functional role of the PrL in AIE-induced social deficits, we inactivated specific neuronal ensembles that were previously activated by a social stimulus. Interestingly, only control males exposed to water during adolescence displayed reduced social investigation following Daun02 inactivation of PrL neuronal ensembles. This finding indicates an important role of the PrL in male social behavior, as others have reported (Amodio & Frith, 2006; Kietzman & Gourley, 2023; Sliwa & Freiwald, 2017). The lack of Daun02 effects on social investigation in AIE-exposed males together with the reduced social behavior following Daun02 inactivation in controls suggests that the neuronal ensembles activated by a social stimulus are likely different between the two exposure conditions. Furthermore, the failure of Daun02 to elicit a change in social behavior of AIE-exposed males, indicates that the PrL ensembles activated in AIE males may not be involved in regulation of social investigation. Given that PrL activation was significantly increased in socially tested animals regardless of sex and adolescent exposure, these ensembles likely are associated with social stimuli, however, due to the complex nature of social behavior, it is possible that these neuronal ensembles regulate different components of social interaction between exposure conditions and sexes. Daun02-induced reduction of social investigation is consistent other studies that used opto- and chemogenetic approaches to reduce activity in the PrL that elicit social deficits (Ferguson & Gao, 2018b; Liu et al., 2020; Przybysz et al., 2023; Scheggia et al., 2020; Xu et al., 2022). For instance, it was previously shown that chemogenetic inhibition of BLA to PrL projections decreased social investigation in control males, whereas males with social deficits induced by prenatal ethanol exposure were insensitive to this manipulation (Przybysz et al., 2023). Importantly, Daun02 inactivated both efferent and afferent PrL connections and although it is possible that social interaction in water- and AIE-exposed males activates different pathways, we are limited in this interpretation. More work is needed to better understand whether different circuits are recruited in AIE vs control males.

In Experiment 2, β-gal expression in the PrL was not different between male adolescent exposure groups that were vehicle-treated, although this difference was evident in Experiment 1. Several reasons may account for this lack of difference in β-gal expression between adolescent exposure conditions, such as disruption of normal PrL function due to cannulation. It is also possible that repeated social testing led to reduced neuronal activation of this region, therefore creating a floor effect in behavior and activation state, likely associated with a decline in novelty of the social test situation. This possibility is supported by findings of reduced social investigation and the number of β-gal+ cells in the PrL of vehicle-treated rats in Experiment 2 compared to our original findings. Furthermore, subsequent social testing has been shown to elicit lower firing rates compared to an initial interaction (Lee et al., 2016).

Although not a direct measure of functional connectivity, β-gal expression was assessed in select downstream ROIs following Daun02 inactivation of the PrL. Both sexes and adolescent exposure groups tended to have lower β-gal expression in limbic structures such as the NAc and CeA. The PrL sends direct projections to the NAc (Cruz et al., 2021), and activation of this pathway has been shown to reduce social preference (Murugan et al., 2017). In contrast, the PrL has been shown to indirectly influence the CeA through the BLA (Gilpin et al., 2015) and paraventricular hypothalamus (PVT) (Kirouac, 2021). However, we did not observe any changes in neuronal activation of the BLA after Daun02 injection; therefore it is possible that the observed decrease in CeA β-gal expression was mediated through the PrL to PVT projection as this pathway has been shown to influence sociability (Yamamuro et al., 2020). Follow up studies should assess whether the PVT is implicated in AIE-induced social deficits. Importantly, Daun02 inactivation decreased social investigation only in control males, with no changes evident in other groups, therefore these possible circuit level alterations in activity likely do not contribute to social behavior.

Our findings of social impairments as well as differential β-gal expression in the IL (home cage) and PrL (home cage and socially tested) in water- and AIE-exposed males were not evident in females. These sex-specific effects of AIE agree with other studies that have found males to be more susceptible to AIE than females (Robinson et al., 2021). In females, neuronal activation of the PrL by interaction with a social partner did not differ between exposure groups. This finding was not surprising, given that AIE-exposed females did not show social deficits. However, as mentioned previously, Daun02 inactivation of PrL neuronal ensembles following social interaction also did not influence social behavior in females, regardless of adolescent exposure. These results are in agreement with the Przybysz et al. (2023) study that reported no changes in female social investigation following manipulation of BLA to PrL inhibition. The regulation of social behavior by the PrL has primarily been assessed in male subjects, however sex differences in social circuitry and underlying neuropeptide systems have been identified (Bredewold & Veenema, 2018; Li et al., 2016). Understanding the neural mechanisms of social behaviors in females is of great importance, and future studies should aim to determine the role of the mPFC in regulating female sociability.

The present interpretations are limited in the absence of knowing which neuronal phenotypes are expressing β-gal and whether our observed effects of inactivation are associated with disrupting afferent or efferent signaling of the PrL. Although the Daun02 inactivation method allows for some specificity in silencing ensembles, targeting cFos expressing cells across the entirety of the PrL eliminates our ability of generating conclusions beyond that of whether the PrL is involved in social behavior. Follow up studies implementing neuronal subpopulation-specific approaches would allow for a more critical understanding of how the PrL is regulating social behavior in water- and AIE-exposed rats.

In conclusion, we identified that AIE exposure increased the basal neuronal activation state of the mPFC, indexed via the cFos proxy β-gal, only in males, but reduced neuronal activation of the CeA in both sexes, irrespective of testing condition. We further found that AIE-induced reductions in social investigation in males was associated with a lower PrL neuronal activation. Using Daun02 pharmacogenetic inactivation of specific PrL neuronal ensembles, we found that the PrL regulates social investigation in control males, with no influence on social investigation of AIE-exposed males and females, regardless of adolescent exposure. Inactivation of neuronal ensembles in the PrL decreased neuronal activation of multiple downstream brain regions, however this was not directly associated with changes to social behavior Additional studies are needed to determine the distinct neuronal phenotypes in the PrL contributing to the observed findings, potentially elucidating a specific mechanism contributing to the aberrant signaling in this region.

## Acknowledgements

Supported by NIH grants U01 AA019972 (Neurobiology of adolescent drinking in adulthood, NADIA), T32 AA025606 (Development and Neuroadaptation in Alcohol and Addictions, DNAA) and F31 AA029300. Any opinions, findings, and conclusions or recommendations expressed in this material are those of the author(s) and do not necessarily reflect the views of the above-stated funding agencies. The authors have no conflicts of interest to declare.

---

**Supplemental Figure 1.** Effects of test conditions on neuronal activation of vOFC, lOFC, NAcC, BLA, and MeA in water- and AIE-exposed males. Significant changes relative to the home cage condition are marked with number signs (# *p* < 0.001); significant changes relative to the context condition are marked with percent signs (% *p* < 0.001).

**Supplemental Figure 2.** Effects of test conditions on neuronal activation of vOFC, lOFC, NAcC, BLA, and MeA in water- and AIE-exposed females. Significant changes relative to the home cage condition are marked with number signs (# *p* < 0.001); significant changes relative to the context condition are marked with percent signs (% *p* < 0.001).

