## Supplementary figures and images for "Determining the neuronal ensembles underlying sex-specific social impairments following adolescent intermittent ethanol exposure"

### Supplemental Figure 1

#### Slide 1
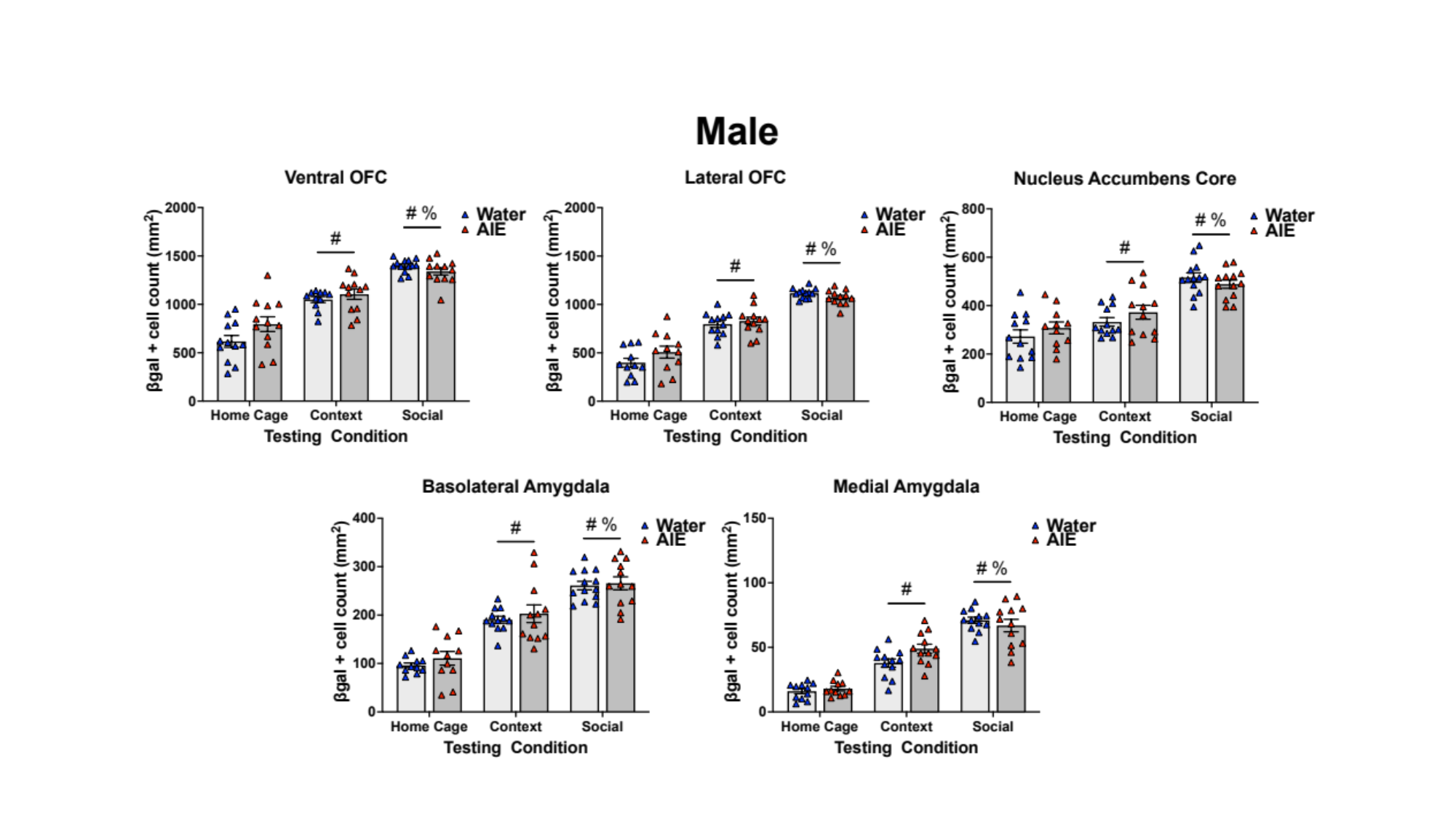

#

### Supplemental Figure 2

#### Slide 1
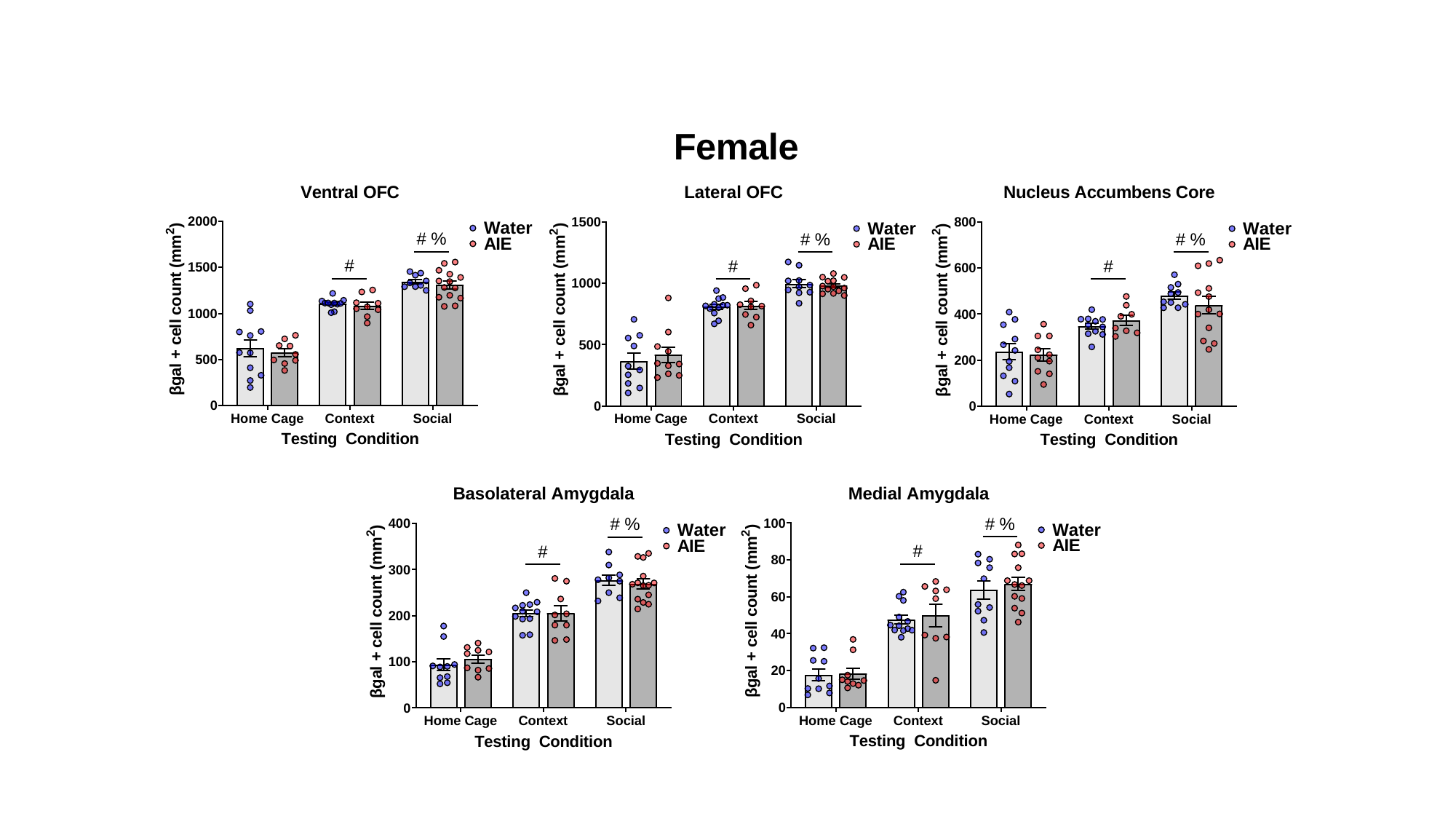
